# *PyPlr*: A versatile, integrated system of hardware and software for researching the human pupillary light reflex

**DOI:** 10.1101/2021.06.02.446731

**Authors:** Joel T. Martin, Joana Pinto, Daniel Bulte, Manuel Spitschan

**Author notes:** Address: Joel T. Martin, Institute of Biomedical Engineering, Department of Engineering Science, University of Oxford, United Kingdom, OX3 7DQ.

## Abstract

We introduce *PyPlr*—a versatile, integrated system of hardware and software to support a broad spectrum of research applications concerning the human pupillary light reflex (PLR). *PyPlr* is a custom Python library for integrating a research-grade video-based eye-tracker system with a light source and streamlining stimulus design, optimisation and delivery, device synchronisation, and extraction, cleaning, and analysis of pupil data. We additionally describe how full-field, homogenous stimulation of the retina can be realised with a low-cost integrating sphere that serves as an alternative to a more complex Maxwellian view setup. Users can integrate their own light source, but we provide full native software support for a high-end, commercial research-grade 10-primary light engine that offers advanced control over the temporal and spectral properties of light stimuli as well as spectral calibration utilities. Here, we describe the hardware and software in detail and demonstrate its capabilities with two example applications: 1) pupillometer-style measurement and parametrisation of the PLR to flashes of white light, and 2) comparing the post-illumination pupil response (PIPR) to flashes of long and short-wavelength light. The system holds promise for researchers who would favour a flexible approach to studying the PLR and the ability to employ a wide range of temporally and spectrally varying stimuli, including simple narrowband stimuli.

## Introduction

The pupillary light reflex (PLR) is the intrinsic mechanism of the pupil to constrict in response to changing light levels. Though its precise biological purpose is still unclear, the PLR is thought to optimise retinal image quality by regulating the amount of light that strikes the retina (Hirata et al., 2003; McDougal & Gamlin, 2015), and it may also help to protect photoreceptors from dangerous levels of light (Laughlin, 1992; Woodhouse & Campbell, 1975). Importantly, as the PLR can be observed directly, it serves as a valuable tool for gaining insight into the integrity and activity of the autonomic nervous system (Girkin, 2003). Indeed, subjective visual assessments of the PLR, such as the swinging flashlight test (Levatin, 1959; Thompson, 1966), are still used routinely in clinical investigations to unmask afferent pupillary defects and give clues to a patient’s neurological state. Though useful in critical care, such techniques are less suited to research due to their limited sensitivity and specificity and the poor inter and intraobserver reliability that exists even among specialists (Litvan et al., 2000; Meeker et al., 2005). The advent and commercial availability of video-based pupillometric techniques in the 1970s enabled researchers and clinical practitioners to make repeatable and precise quantitative pupil measurements. Consequently, the pupil’s response to light is now well characterised in both health and disease (Loewenfeld, 1993).

The aperture of the pupil at any given time depends on the tone of the *dilator* and *sphincter pupillae*—the two opponent smooth muscles of the iris. The iris sphincter receives parasympathetic innervation and is almost solely responsible for the constriction of the pupil that follows an increase in retinal illumination (McDougal & Gamlin, 2015). When light strikes the retina, photons are absorbed by photoreceptors and the neural signal traverses a short reflex arc comprising the photoreceptor, bipolar and ganglion cells of the retina (as well as other interneurons), the olivary pretectal nucleus of the midbrain and the Edinger-Westphal nucleus, which projects to the iris sphincter muscle via the ciliary ganglion (Hall & Chilcott, 2018). Following a sudden flash of white light, a normal pupil will begin to constrict after approximately 230 ms and, after reaching peak constriction, will enter a redilation phase and return to baseline. Redilation of the pupil upon light cessation depends on two integrated processes: relaxation of the sphincter muscle due to parasympathetic inhibition and contraction of the dilator muscle following excitation in the sympathetic pathway (Szabadi, 2018). The PLR is typically parametrised in terms of the latency, amplitude, velocity and acceleration of change in pupil size and its dynamics are affected by normal ageing (Bitsios et al., 1996; Winston et al., 2019). In a broad range of ophthalmic, neurologic, and psychiatric conditions (Chen et al., 2011; Girkin, 2003; Van Stavern et al., 2019), the PLR can be abnormal, making it an important tool in research and diagnostics (Hall & Chilcott, 2018; Troiani, 2020).

Where it was once assumed that the PLR is controlled entirely by the integration of signals from rod and cone photoreceptors, we now know that steady-state pupil size is largely under the influence of intrinsically photosensitive retinal ganglion cells (ipRGCs)—a subpopulation of retinal ganglion cells which express the photopigment melanopsin in their axons and soma (Clarke et al., 2003a; Provencio et al., 2000). ipRGCs are sensitive to high intensity, short-wavelength (blue) light and control non-visual functions, such as circadian photoentrainment and pupil size (Spitschan, 2019), via direct projections to the suprachiasmatic nucleus of the hypothalamus and the olivary pretectal nucleus (Do, 2019), respectively. The post-illumination pupil response (PIPR) describes the sustained constriction of the pupil following exposure to short-wavelength light, usually relative to long-wavelength light, and is assumed to be a unique non-invasive biomarker of melanopsin function in the human retina (Adhikari et al., 2015; Clarke et al., 2003b; Kankipati et al., 2010). Like the flash response to white light, the PIPR is researched extensively for its potential as a biomarker in various ocular and neurodegenerative diseases (Chougule et al., 2019; Feigl & Zele, 2014; Kankipati et al., 2011).

Researching the PLR requires a system for illuminating the retina and measuring pupil size simultaneously. For patient monitoring in critical care, hand-held pupillometers offer an attractive all-in-one solution as they are portable, reliable and easy to use (Meeker et al., 2005; Taylor et al., 2003). These ‘point-and-shoot’ devices are aimed at the eye to deliver a light stimulus and use infrared illumination, video recording and internal algorithms to provide an instantaneous readout of PLR parameters. Some limitations of automated pupillometers which make them less suited for scientific research are that they can be expensive and inflexible, offering minimal control over stimulus parameters (e.g., duration, wavelength, intensity) and in some cases no access to the raw data. Conversely, video-based eye trackers, which usually measure pupil diameter or area as part of their gaze estimation pipeline, are often favoured in research for their versatility. But video-based eye trackers and similar recording devices must be integrated with a system for administering light stimuli. This task may not prove too challenging for basic experiments where a standard computer screen will suffice, but it becomes more challenging when research calls for a bespoke setup to control the spatial extent of retinal stimulation and the spectral and temporal properties of light stimuli. One solution is to use a Maxwellian view pupillometry system (e.g., Adhikari et al., 2015; Cao et al., 2015; Kankipati et al., 2010; Westheimer, 1966), where the light stimulus is focused onto an aperture placed in front of the eye, or in the entrance plane of a pharmacologically dilated pupil, and the consensual pupil response is measured from the other eye. An alternative, which does not require complex optical engineering, pharmacological dilation of the pupil, or strict fixation control on the part of the participant, is to use a full-field—‘Ganzfeld’—illumination system (e.g., Bonmati-Carrion et al., 2018; Kardon et al., 2009); however, commercial solutions for this mode of stimulation can be prohibitively expensive.

Here we describe *PyPlr* (Martin & Spitschan, 2021)—a custom Python software that works with the Pupil Core (Pupil Labs GmbH, Berlin, Germany) eye-tracking platform to offer an affordable, versatile, extensible and transparent solution for researching the PLR. Features include: 1) user-friendly and feature-rich interfaces to Pupil Core (Pupil Labs, GmbH, Berlin, Germany), Spectra Tune Lab (STLAB: LEDMOTIVE Technologies, LLC, Barcelona, Spain) light engine and Ocean Optics (Ocean Insight Inc., Oxford, UK) spectrometers, 2) flexible support for alternative stimulus delivery and measurement systems, and 3) scripting tools to facilitate stimulus design, optimisation and delivery, communication with respect to timing, and extraction, cleaning, and analysis of pupil data. We also describe how full-field, homogenous stimulation of the retina can be achieved with a low-cost integrating sphere that serves as an alternative to the more-complex Maxwellian view pupillometry setup. Following a detailed overview of the hardware and the software we present two example applications as a proof of concept: 1) pupillometer-style measurement and parametrisation of the PLR to a flash of white light, and 2) measuring the post-illumination pupil response (PIPR) to flashes of long vs. short-wavelength light.

## Overview

*PyPlr* is an open-source Python software for researching the PLR with the Pupil Core eye-tracking platform. The software, which is mapped out graphically in Figure 1, comprises a set of modules for interfacing with hardware, obtaining measurements, designing and running experimental protocols, and processing pupil data. The project is maintained on GitHub (https://github.com/PyPlr/cvd_pupillometry) under the MIT License with extensive documentation (https://pyplr.github.io/cvd_pupillometry/) and registered with the Python Package Index (https://pypi.org/project/pyplr/) making it installable via the packaging tool *pip*.

**Figure 1.**
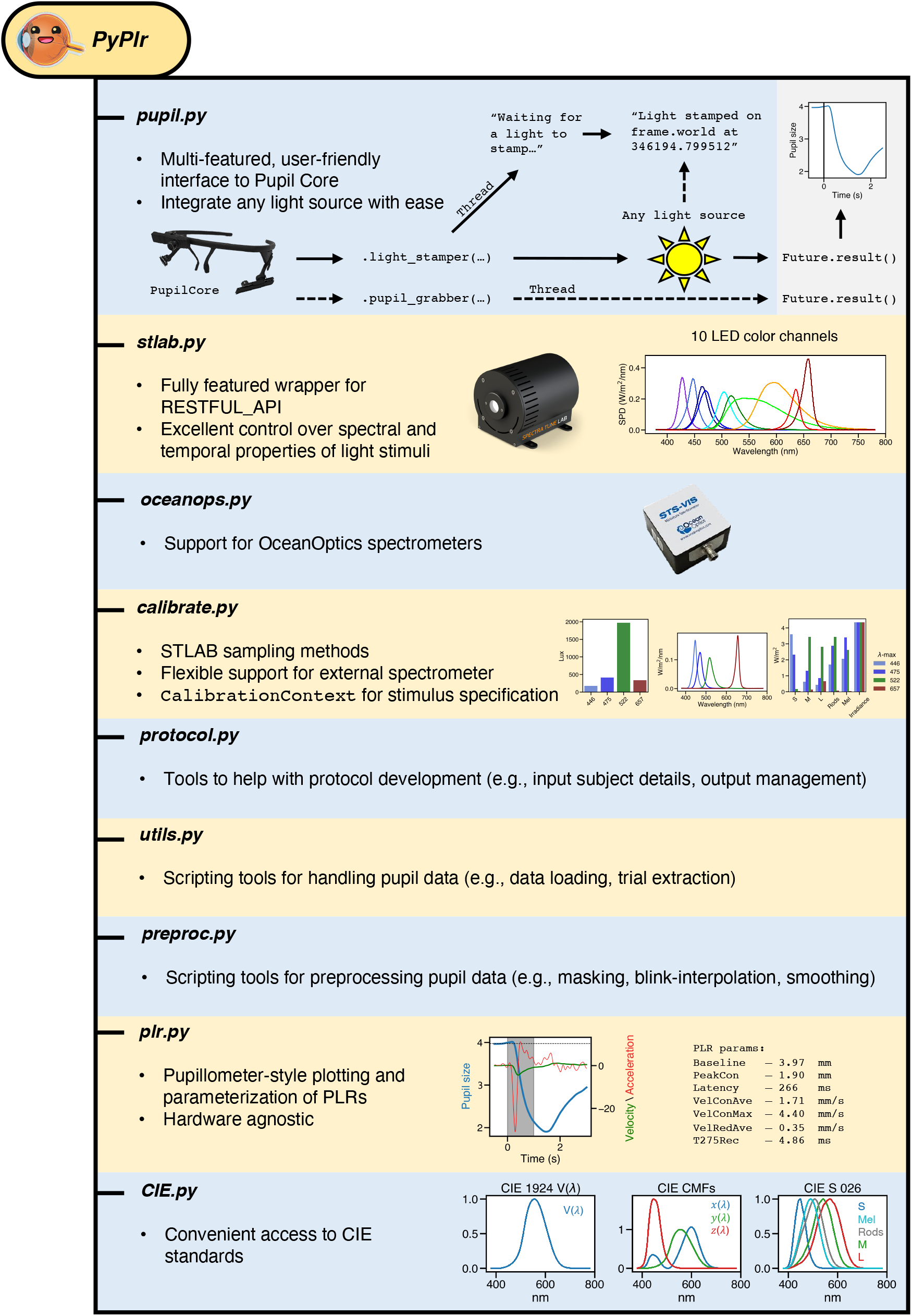
*PyPlr* software overview.

A key feature of *PyPlr* is that light stimuli can be timestamped with good accuracy using the Pupil Core World camera. This feature makes it easy to integrate any light source given a suitable geometry. For our own stimulation and measurement system we developed a low-cost integrating sphere (see Figure 2 and description below) for use with STLAB, but *PyPlr*’s native support for timestamping opens the door to alternative solutions. In this section we present an overview of the key features of *PyPlr* and describe the low-cost integrating sphere that we built for our stimulation and measurement system.

**Figure 2.**
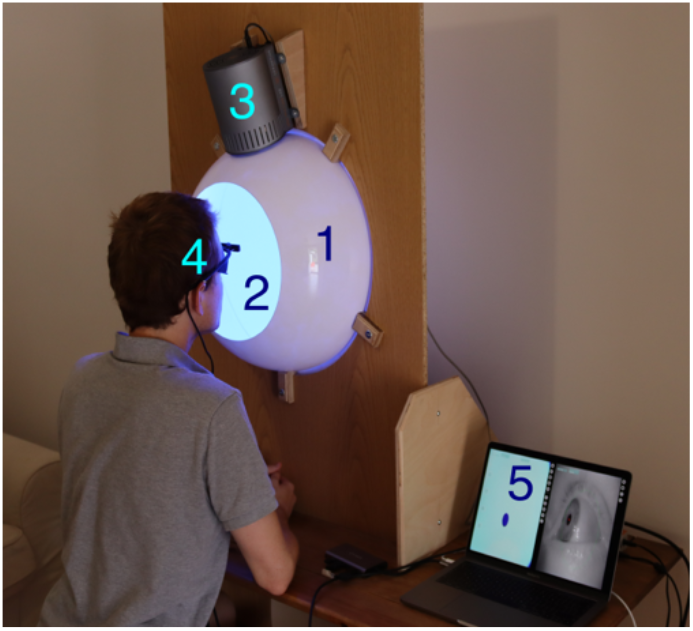
Stimulation and measurement system: 1) integrating sphere constructed from two acrylic half-domes, housed and stabilized with a wooden fixing plate, 2) inside coating of Avian-B high reflectance paint to scatter light homogenously, 3) STLAB light source mounted above entry port, 4) Pupil Core eye-tracking headset, and 5) laptop running Pupil Capture and custom Python software. The photograph was taken with the participant’s permission.

### *PyPlr* and Pupil Core

*PyPlr* works with Pupil Core—an affordable, open-source, versatile, research-grade eye-tracking platform with high sampling rates, precise model-based 3D estimation of pupil size, and many other features which make it well-suited to our application (see Kassner et al., 2014, for a detailed overview of the system). Of note, Pupil Core has a Network API which supports fast and reliable communication and real-time access to data via *ZeroMQ*, a universal messaging library, and *MessagePack*, a binary format for information interchange. As noted above, *PyPlr* leverages the real-time data streaming capabilities of Pupil Core’s forward-facing World camera to timestamp the onset of light stimuli with good temporal accuracy, opening the door to integration with virtually any light source given a suitable geometry. A Pupil Core headset and its accompanying software (i.e., *Pupil Capture*) is therefore a basic dependency of a functioning *PyPlr* setup.

#### pyplr.pupil

*PyPlr*’s *pupil.py* module greatly simplifies working with Pupil Core and its Network API by wrapping all of the tricky *ZeroMQ* and *MessagePack* code into a single device class. The *PupilCore* device class has a *.command(…)* method giving convenient access to all of the commands available via *pupil remote*, which makes it trivially easy to connect to the eye tracker and perform basic operations, such as starting and stopping a recording, calibrating, getting the current pupil time, and so forth. *PupilCore* also has a rich set of class methods to facilitate the design and implementation of effective pupillometry protocols. Readers are encouraged to refer to the code and online documentation for detailed information on the full range of functionality. Here we describe two key methods—*.light_stamper(…)* and *.pupil_grabber(…)*—and the problems they were designed to solve. A minimal example of how to use *PupilCore* and its class methods to measure and plot a PLR to any light stimulus is provided in Figure 3.

**Figure 3.**
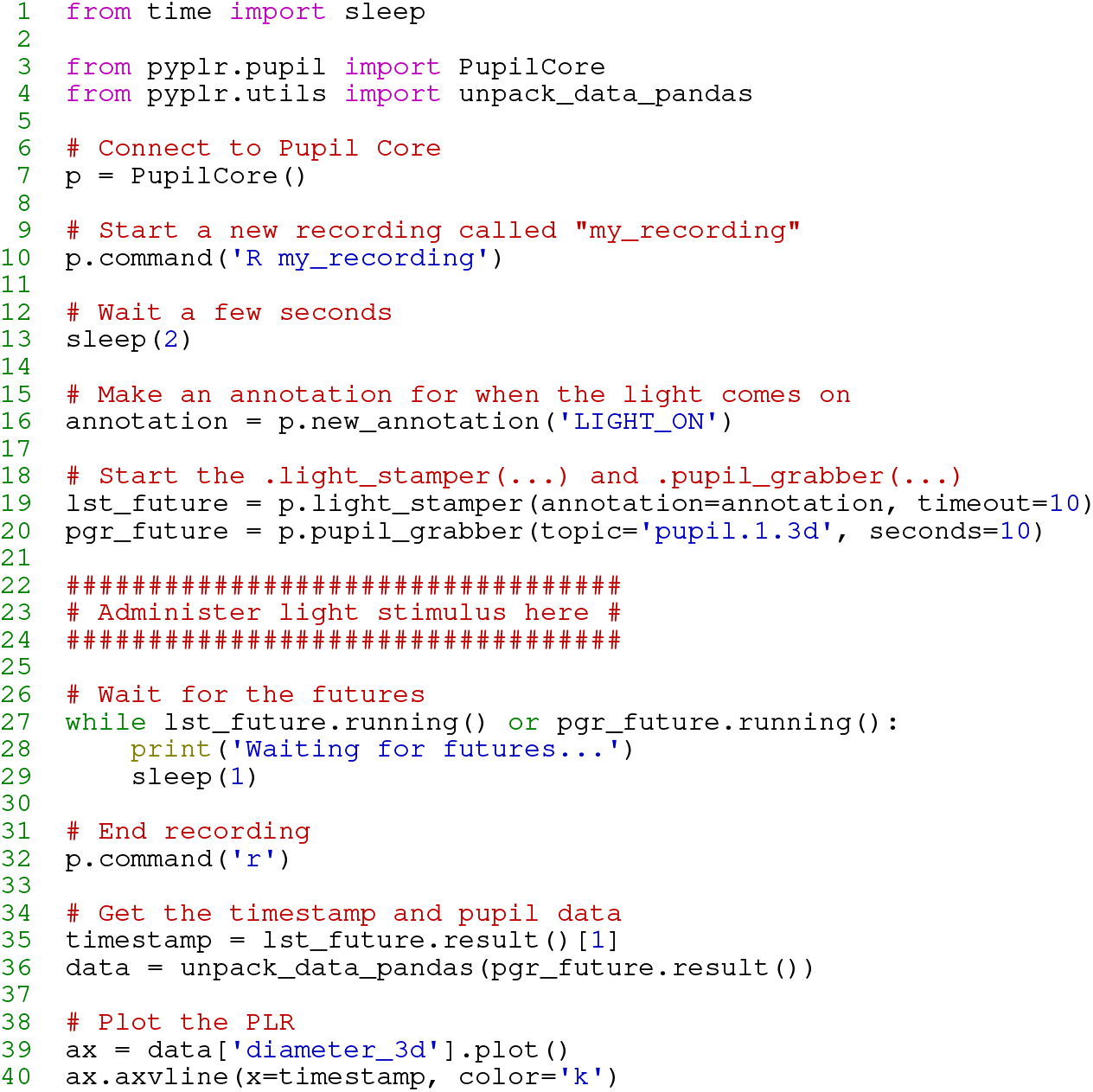
Minimal example demonstrating the use of the *PupilCore* device class and its *.light_stamper(…)* and *.pupil_grabber(…)* methods for real-time PLR measurement. Note that it is not necessary to make a recording for these methods to work, and that the script will work for any light stimulus that can be detected by the World camera (e.g., a computer screen, a light switch in a dark room, an integrating sphere).

*.light_stamper(…).* To extract experimental events and calculate time-critical PLR parameters (e.g., constriction latency, time-to-peak constriction) requires a reliable indication in the pupil data of the time at which a light stimulus was administered. The Pupil Capture software has an *Annotation Capture* plugin which allows for samples to be labelled with an annotation manually via keypress or programmatically via the Network API in a process that is analogous to sending a ‘trigger’ or ‘event marker’. The obvious way to timestamp a light stimulus therefore would be to control the light source programmatically from a Python script and send an annotation immediately before or after issuing a command to the light; but, as a universal approach, this will likely prove far from ideal, because different light sources have their own latencies which are often variable and difficult to reference. In fact, our own light source (described below) takes commands via generic HTTP requests and has a variable response time on the order of a few hundred milliseconds. Given that we may want to calculate latency to the onset of pupil constriction after a temporally precise light stimulus, such variability is unacceptable.

To solve the timestamping issue in a way that makes it easy to integrate *PyPlr* and Pupil Core with any light source, we developed the *.light_stamper(…)*—a *PupilCore* class method that uses real-time data from the forward facing World camera to timestamp the onset of a light stimulus based on a sudden change in the average RGB value. The underlying algorithm simply keeps track of the two most recent frames from the World camera and sends an annotation with the timestamp of the first frame where the average RGB difference exceeds a given threshold. Crucially, a *.light_stamper(…)* runs in its own thread with Python’s *concurrent.futures*, so the flow of execution is not blocked and the result—i.e., the timestamp—is available via a call to the *.result()* method of a returned *Future* object once the light has been stamped. To work properly, the *.light_stamper(…)* requires a suitable stimulus geometry (the camera must be able to see the light source), an appropriately tuned threshold value, and the following settings in Pupil Capture:

1. *Auto Exposure Mode* of the camera must be set to *Manual*
2. *Frame Publisher Format* must be set to *BGR*
3. *Annotation Capture* plugin must be enabled

With our integrating sphere setup, we find that the *.light_stamper(…)* flawlessly captures the first frame in which a light stimulus becomes visible for a range of practical intensities, as verified using Pupil Player and the Annotation Player plugin. Timestamping accuracy, therefore, is limited only by camera settings (e.g., frame rate) and how well the Pupil software can synchronise the clocks of the Eye and World cameras. We were able to test camera clock synchronisation by putting the Pupil Core headset inside our integrating sphere (described below) and repeatedly flashing a bright orange light containing enough near-infrared to afford detection by the Eye cameras as well as the World camera. Before each flash, concurrent *.light_stamper(…)*’s were instantiated, giving us the timestamp of the frame where the luminance change was detected independently for each camera. Knowing from community discussions that the Pupil software handles timestamps differently on Windows and Unix operating systems, and more generally that frame rate will play an important role in determining the accuracy of the *.light_stamper(…)*, we performed the test (*n* = 100 light flashes) on both macOS (Big Sur, 11.3.1) and Windows (Windows 10) with frame rates of 60 and 120 for all cameras (Pupil Capture v3.2-20). For each run of the protocol, Eye camera resolution was kept at (192, 192) with Absolute Exposure Time of 25, and for the World camera, (640, 480) and 60. Auto Exposure Mode was set to ‘manual mode’ for all cameras, and Auto Exposure Priority was disabled for the World camera.

The effect of frame rate and operating system on timestamping is shown in Figure 4. For both macOS and Windows, the Eye camera timestamps appear well-synchronised with a margin of error that is to be expected given the frame rate. On Windows, the World camera timestamps fell consistently around 60 ms before the Eye camera timestamps at both 60 and 120 FPS. The same pattern of a leading World timestamp was observed, though to a lesser degree, with macOS. The timestamps appeared best synchronised overall on macOS with cameras running at 120 FPS, where the World camera led by 15 ms on average.

**Figure 4.**
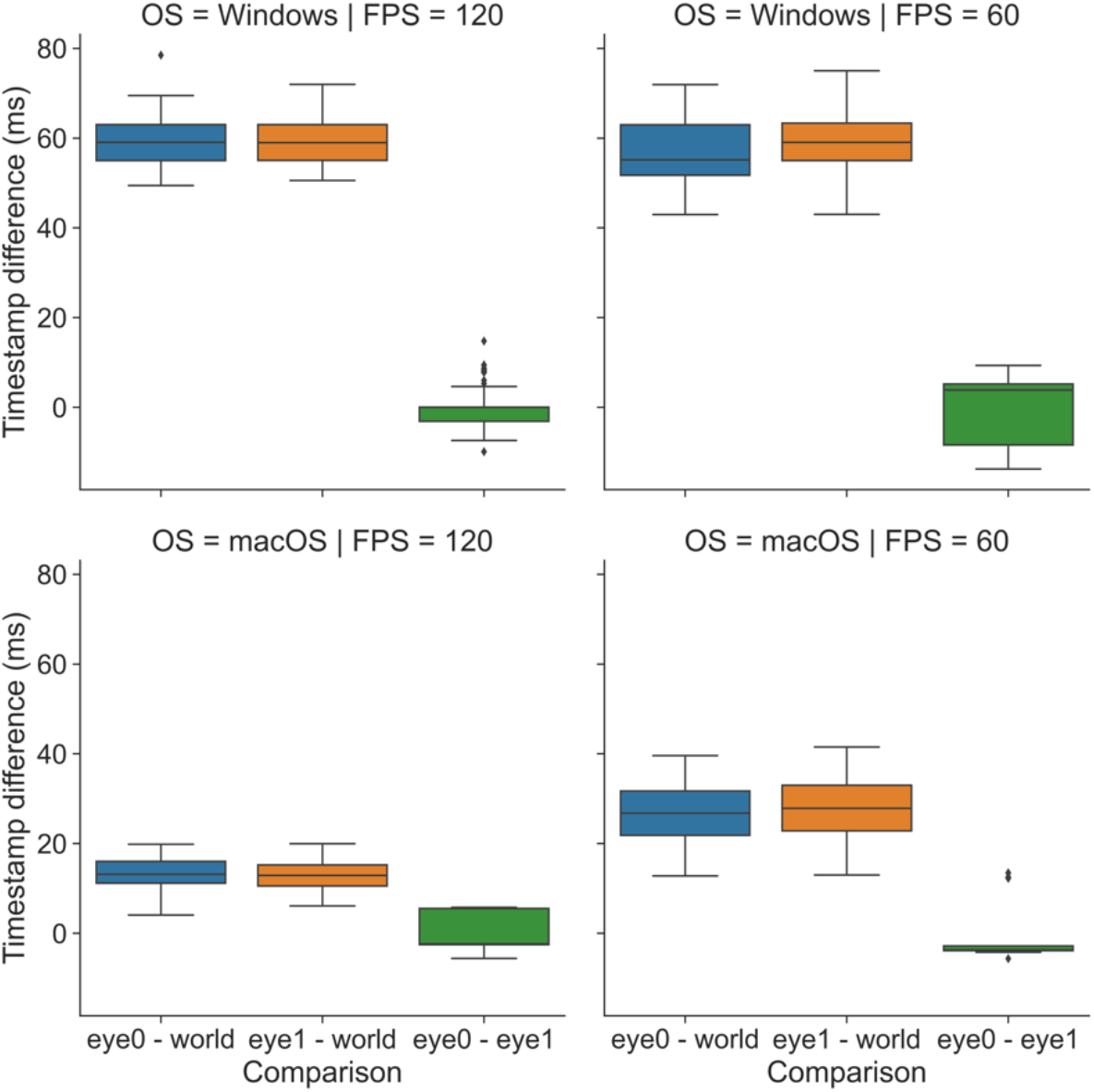
Effect of operating system (OS: macOS vs. Windows) and frame rate (FPS: 60 vs. 120) on timestamp differences for light flashes (*n* = 100) detected independently for each Pupil Core camera with concurrent *.light_stamper(…)*’s.

Understanding what underlies these discrepancies requires a developer’s knowledge of the Pupil software and its treatment of timestamps on different operating systems. At the time of writing, we understand from community discussions that macOS and Linux use the hardware timestamps generated by the cameras at the start of frame exposure, whereas Windows uses software timestamps generated by *pyuvc* using the system’s monotonic clock at the time when the frame is done transferring from camera to computer. Unlike hardware timestamps, the Windows software timestamps are subsequently corrected by subtracting a fixed amount of time corresponding to the approximate camera latency (i.e., the difference between software and hardware timestamps), but at present this procedure assumes the default resolution of the camera in question and is not optimised to account for the different camera latencies associated with different resolutions (N.B., larger frames take longer to transfer). This may be optimised in a future update to the Pupil software. At present, the implication for our application is as follows: time-critical measures of a PLR referenced to a World camera *.light_stamper(…)* timestamp will be consistently overestimated by 15 to 60 ms, depending on the operating system and camera settings being used. Though not ideal, the timestamp discrepancy is at least repeatable and potentially correctable, meaning researchers are free to obtain time-critical measurements of the PLR. For applications that require precise timing, researchers should perform their own due diligence and engage in discussions with the Pupil Labs community to better understand the timestamping implementation of the Pupil software.

*.pupil_grabber(…).* The *.pupil_grabber(…)* is a *PupilCore* class method that simplifies real-time access to data and empowers users to design lean applications that bypass the sometimes-cumbersome record-load-export routine of the Pupil Player software. As arguments, the *.pupil_grabber(…)* takes a topic string specifying the data to be grabbed (e.g., *pupil.1.3d* to grab 3D model data for the left eye, *pupil.* to grab all pupil data, etc.) and a numerical value specifying the number of seconds to spend grabbing data. Like the *.light_stamper(…)*, the *.pupil_grabber(…)* runs in its own thread with *concurrent.futures* and gives access to data via a call to the *.result()* method of a returned *Future* object after the work is done. Grabbed data are stored as a list of dictionaries and can subsequently be organised into a more manageable format with the *unpack_data_pandas(…)* helper function from *pyplr.utils.*

### Spectra Tune Lab light source

As a light source for our stimulation system we chose Spectra Tune Lab (STLAB: LEDMOTIVE technologies LLC, Barcelona, Spain)—a high-end, spectrally tuneable light engine with ten LED colour channels, capable of generating a broad range of spectral compositions. The gamut of the device and the spectral power distributions for each LED channel at maximum are displayed in Figure 5 and Figure 6, respectively. STLAB connects via network cable to a small computer called the Light Hub (a Beaglebone board running Linux), which connects to a controlling computer via USB or some network protocol (e.g., LAN, WAN, internet, etc.). STLAB can be controlled programmatically with most languages via its REST API, which works with generic GET and SET operations. Spectra are most easily defined by passing an array of ten 12-bit integers to set the intensity of each individual LED channel. Here we describe *pyplr.stlab*, *PyPlr*’s module for interfacing with STLAB, and review key aspects of performance and functionality.

**Figure 5.**
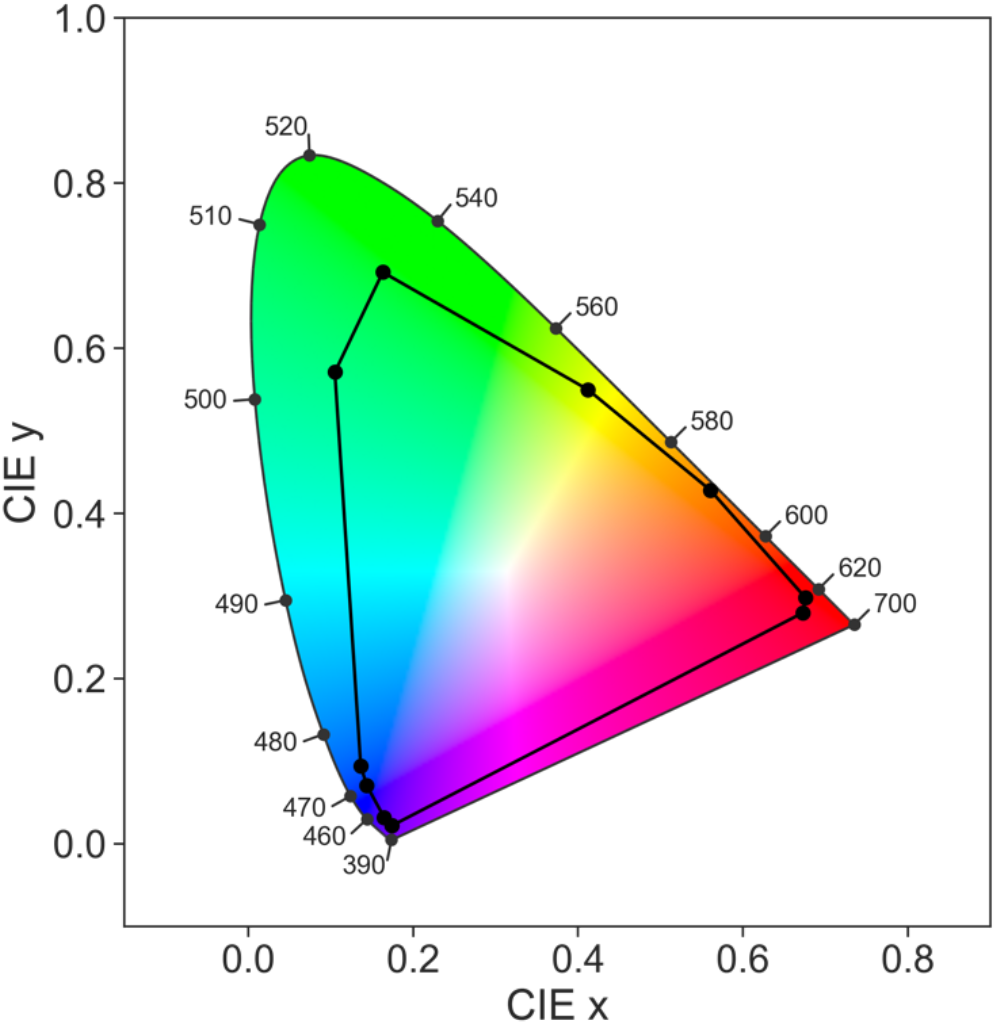
CIE 1931 ‘horseshoe’ chromaticity diagram (2° standard observer) for STLAB’s ten LED channels at maximum, defining the gamut of the stimulation system. Spectral data were obtained in a darkened room with an OceanOptics STS-VIS (Ocean Insight Inc., Oxford, UK) spectrometer at the plane of the integrating sphere viewing port.

**Figure 6.**
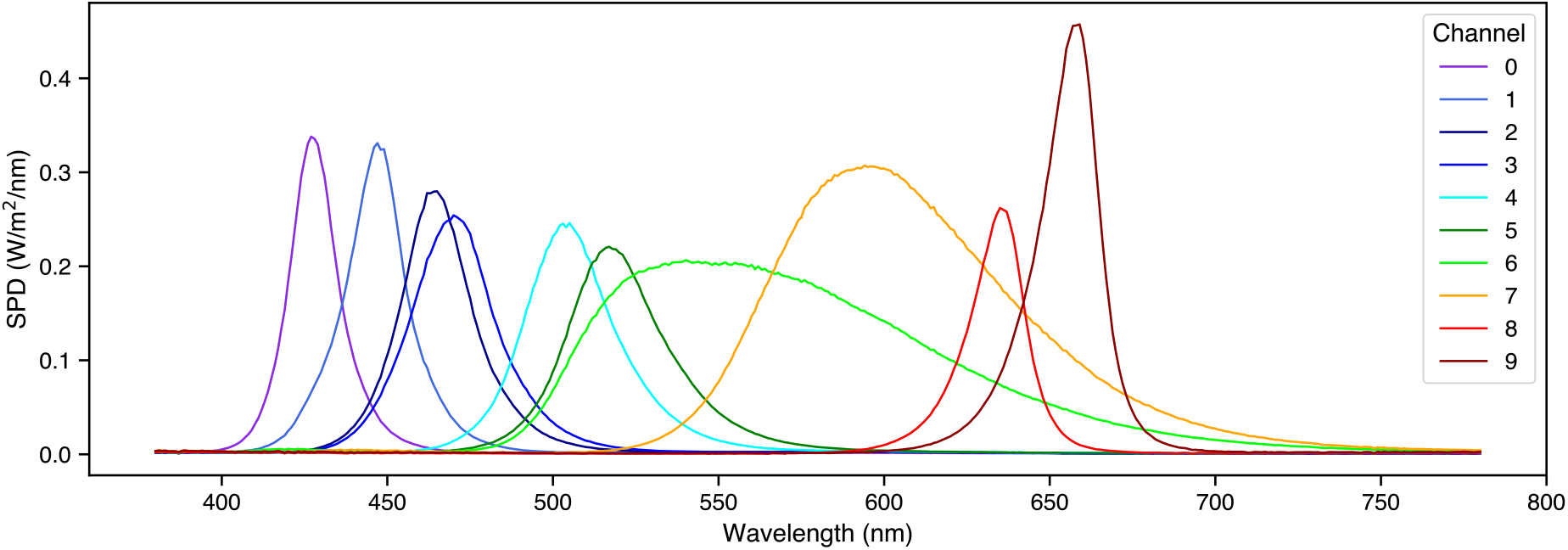
Spectral power distributions for STLAB’s ten LED channels at maximum. Spectral data were obtained in a darkened room with an OceanOptics STS-VIS (Ocean Insight Inc., Oxford, UK) spectrometer at the plane of the integrating sphere viewing port.

#### pyplr.stlab

This module contains *SpectraTuneLab*, a device class that uses the Python *requests* library to wrap all of the functions from STLAB’s REST API. Readers are encouraged to check the code and documentation for further information. Additional helper functions are included to assist with developing stimuli. Note that a license is required to develop against the REST API.

#### Device timing and video files

STLAB operates synchronously by default, meaning that all commands sent by the Light Hub must be acknowledged before a new instruction can be processed. According to the device manual, response times in this mode of operation are on the order of around 250 milliseconds. We verify this with our own testing, but also note that on rare occasions, perhaps when the Light Hub is busy processing other tasks, the response time can be up to five s. Such a delay is not suitable for administering light stimuli requiring exact timing. To do this, we leverage STLABs asynchronous mode of operation, which allows for real-time spectral streaming with a spectral switching time of less than ten milliseconds (i.e., one spectrum every ten milliseconds). This mode of operation requires the advanced preparation of *video files*, which are JSON files with a particular structure and the idiosyncratic DSF—*dynamic sequence file*—extension. The core inputs for making a video file are a time vector to specify the spectral switching times and a separate list of spectra (specified as arrays of ten 12-bit integers). *pyplr.stlab* has a *make_video_file(…)* function which will convert an appropriately structured *pandas* (McKinney, 2010) *DataFrame* into the required JSON format and save it with a DSF extension. Also included are some higher-level convenience functions for quick and easy specification of timed pulse stimuli. To use video files in an experimental protocol, one must simply use the *.load_video_file(…)* and *.play_video_file(…)* methods of the *SpectraTuneLab* device class.

### Integrating sphere

For some experiments it may be sufficient to perform light stimulation with a standard computer monitor, but where research calls for advanced control over the geometry of retinal stimulation, a bespoke setup is needed. One solution is to use a Maxwellian view pupillometry system, where the light stimulus is focused onto an aperture positioned in front of the eye or in the entrance plane of a pharmacologically dilated pupil, and the consensual pupil response is measured from the other eye (e.g., Cao et al., 2015). But this approach requires optical engineering and resources that may not be available in the average research setting. As an alternative, we developed a low-cost integrating sphere (Figure 2) that delivers a full-field, ‘Ganzfeld’ stimulus and precludes the need for optical engineering, pharmacological dilation of the pupil, and strict fixation control on behalf of the participant.

#### Construction

We built the sphere from two 45-cm diameter flanged acrylic half-domes (Project Plastics Ltd., Colchester, UK). The inside surfaces of the domes were cleaned, keyed with a scotch pad and then primed with Zinsser B-I-N Off white Matt Primer & undercoat Spray paint (William Zinsser & Co. Inc., Birtley, UK) before they were sprayed with multiple coats of Avian-B high reflectance paint (Avian Technologies LLC, New London, NH). The Avian-B premix was mixed on a magnetic mixing plate with the correct quantities of denatured alcohol and distilled water and tested for viscosity and pH in accordance with the application notes. A 28 cm opening in one of the domes serves as a viewing port, and an additional 7 cm opening (subtending ~9° from the plane of the viewing port) opposite the viewing port was included to allow for secondary stimuli (e.g., a fixation target) or to afford exclusion of the foveal macular pigment from stimulation. On the same half of the sphere as the viewing port, a 30 mm entry port for the STLAB light source was cut at an angle of 22.5-deg from the top such that the diffuser of the light source could not be seen directly when looking straight ahead. The sphere was stabilized on a wooden fixing plate making it suitable for placement on a desk and for use with a chinrest. The raw materials for the integrating sphere cost us less than £1500.

#### Calibration

To create a calibrated forward model of the STLAB-sphere rig that represents what an observer actually sees when looking into it, we obtained measurements with an external spectrometer positioned at the plane of the viewing port. The *pyplr.calibrate* module was designed to streamline this process with a *SpectraTuneLabSampler(…)* class—a sub-class of *pyplr.stlab.SpectraTuneLab* with added sampling methods and support for an external spectrometer. Any spectrometer with a python interface can be integrated here with minimal effort, but we used an Ocean Optics STS-VIS (Ocean Insight Inc., Oxford, UK), which has native support from *pyplr.oceanops* via the *Seabreeze* (v1.3.0; Poehlmann, 2019) Python library. It would take a long time to sample every possible device setting, so we opted to sample the 12-bit intensity range in a dark room, independently for each LED channel in steps of 65, which amounts to 63 evenly spaced measurements per LED. Figure 7 shows how easy it was to obtain these spectral data.

**Figure 7.**
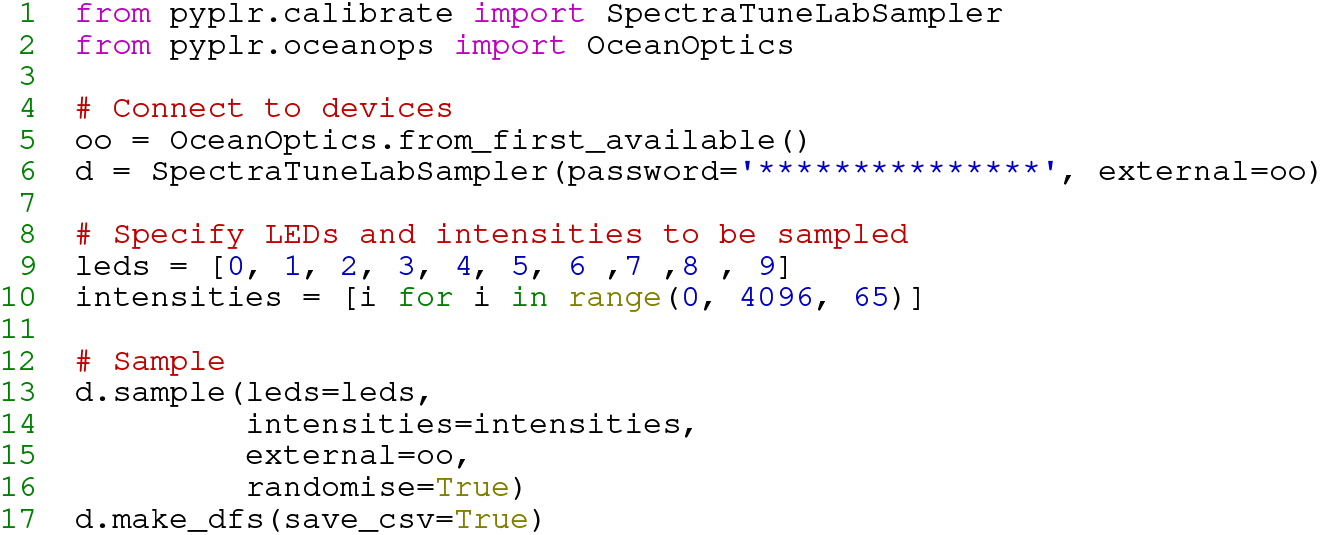
Profiling the integrating sphere with *pyplr.calibrate* and *pyplr.oceanops*. Measurements were obtained in a dark room with an OceanOptics STS-VIS (Ocean Insight Inc., Oxford, UK) spectrometer fitted with a cosine corrector and positioned at typical eye position.

Having obtained the raw spectral measurements with our OceanOptics spectrometer, a device-specific calibration pipeline was implemented to account for the effect of PCB temperature and integration time on raw sensor readings. The calibrated spectral data were then passed to *pyplr.calibrate.CalibrationContext*, a data-handling class which uses reindexing and linear interpolation to fill in the gaps and automatically generate lookup tables giving easy access to the predicted spectral power distribution, *a*-opic irradiances, lux, and unweighted irradiance for all possible combinations of LED-intensity settings. Crucially, the *CalibrationContext* also has a *.predict_spd(…)* method that will predict the spectral output from a list of ten 12-bit values, as required by STLAB. There is also a *.fit_curves(…)* method that fits beta cumulative distribution functions to the LED-intensity data, and an *.optimise(…)* method that applies the resulting parameters to correct a stimulus profile for any departures from linearity. Figure 8 shows how spectra can be accurately predicted from the *CalibrationContext* and Figure 9 demonstrates the linearity of the relationship between STLAB’s input and output.

**Figure 8.**
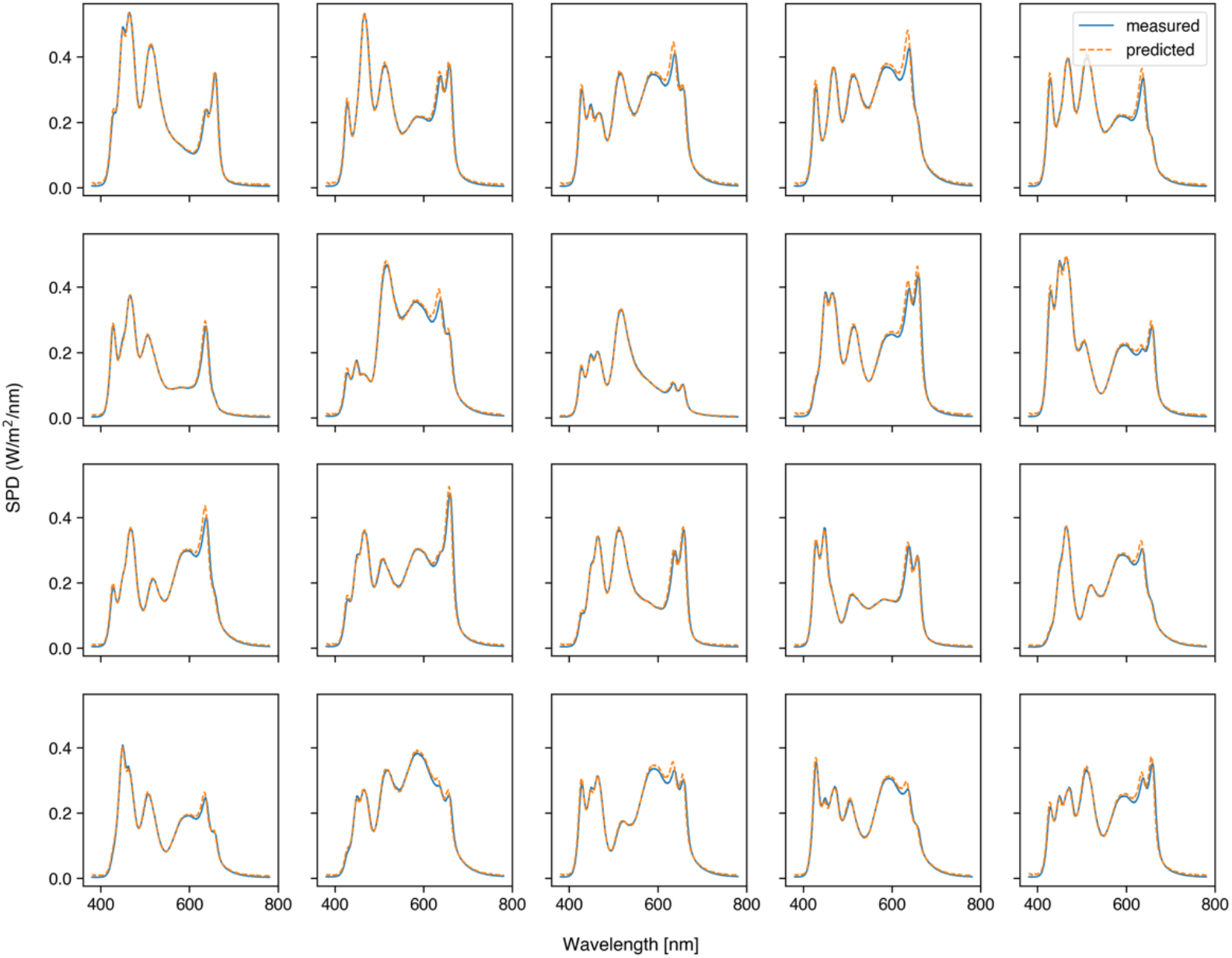
Measured spectral power distributions for 20 random device settings compared with the spectral power distributions as predicted by the *CalibrationContext.predict_spd(…)* method using the same settings. The 20 random spectra were measured with the same spectrometer and under the same conditions as the calibration spectra.

**Figure 9.**
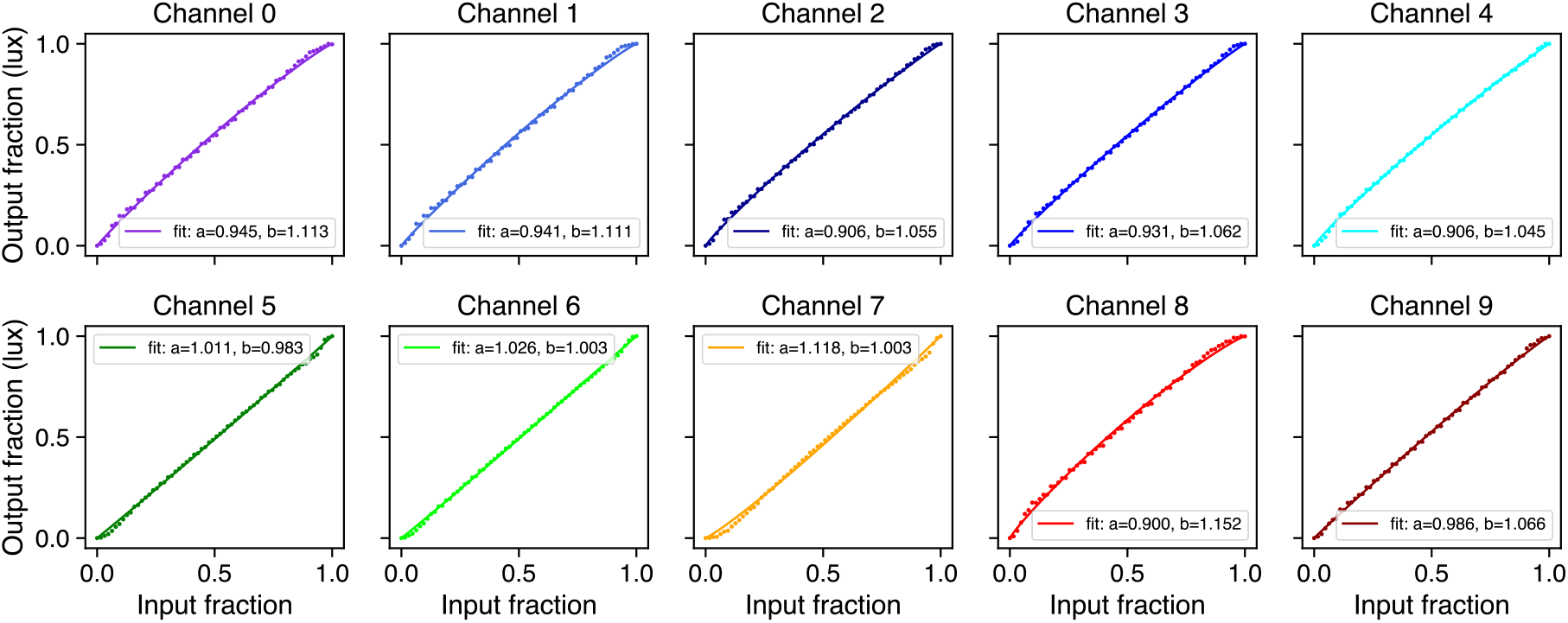
Output of the *CalibrationContext.fit_curves(…)* method, showing the relationship between input (12-bit) and output (photopic illuminance in lx) for all of STLAB’s LED channels, as measured by an OceanOptics STS-VIS (Ocean Insight Inc., Oxford, UK).

#### Safety

We evaluated the safety of the stimulation system in accordance with the British Standards Document on the Photobiological Safety of Lamps and Lamp Systems (BS EN 62471: British Standards Institute, 2008). Section 4.1 (Annex ZB, page 40) of the BS EN 62471 states that ‘Detailed spectral data is required if the luminance of the source exceeds 10^4^ cd/m^−2^’. Initial scoping measurements collected with a Photo Research SpectraScan PR-670 for all LEDs at 100% gave a luminance reading of 18000 cd/m^2^ at the plane of the viewing port. The maximum output of our stimulation system therefore exceeded this specification, so we obtained detailed spectral measurements. Section 4.3.3 of the BS EN 62471 states:

To protect against retinal photochemical injury from chronic blue-light exposure, the integrated spectral radiance of the light source weighted against the blue-light hazard function, *B*(λ), i.e., the blue light weighted radiance, LB, shall not exceed the levels defined by:

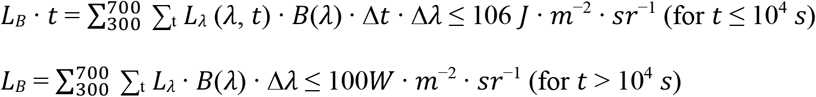

Where:

*L*_λ_ (λ, *t*) is the spectral radiance in *W* · *m*^−2^ · *sr*^−1^ · *nm*^−1^

*B*(λ) is the blue-light hazard weighting function

Δλ is the bandwidth in nm

*t* is the exposure duration in seconds. (p. 44)

Using the minimum radiance limit for the retinal blue light hazard exposure limit, given as 100 *W* · *m*^−2^ · *sr*^−1^ for exposures of greater than 10000 s, we note that our source is below the retinal blue light hazard exposure limit. These findings were confirmed by processing the data with “EyeLight”, an Optical Safety Software Platform supplied by Blueside Photonics Ltd. (Preston, UK) and the National Physical Laboratory (Teddington, UK). However, given that our stimulation system may be used in a dark room following a period of dark adaptation, pupil diameter will be greater than 3 mm at the start of exposure. Section 4.2.1 of the BS EN 62471 states:

When the luminance of the source is adequately high (>10 cd.m^−2^), and the exposure duration is greater the 0.25s, a 3mm pupil diameter (7mm^2^ area) was used to derive the exposure limit. (Annex ZB, p. 40)

To take this into account we applied a pupil correction factor of 6 (pupil ratio: 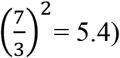 which reduces the retinal blue light hazard exposure limit to 16.6 *W* · *m*^2^ · *sr*^−1^. Therefore, when running the source at 100% and applying a safety factor to correct for the pupil size, our stimulation system is above the radiance retinal blue light hazard exposure limit value of 100 *W* · *m*^2^ · *sr*^−1^ for an exposure of 10000 s. Considering, however, that the PLR is a component of the aversion response to bright light under normal viewing conditions and that we are only presenting 1 second pulses of light, we conclude from this analysis that our system is safe for our intents and purposes. For protocols involving prolonged exposure to short wavelengths or pharmacological pupil dilation, researchers should consider the safety implications and consult the relevant standards to ensure that stimuli do not exceed the retinal blue light hazard exposure limit.

### Data analysis

There is more than one valid approach to the analysis of pupillometry data, but the optimal approach will depend on the type of experiment being run, the quality of the data, and the research question in mind. Kelbsch et al. (2019) give an informative view on standards in pupillometry of the light reflex and many papers offer advice on best practices and specific issues to do with data analysis (e.g., Hayes & Petrov, 2015; Kret & Sjak-Shie, 2019; Mathôt, 2017; Sirois & Brisson, 2014; Winn et al., 2018), much of which is embodied in community-developed packages that aim to streamline the processing and analysis of pupillometry data (e.g., Acland & Braver, 2014; Mittner, 2020). Ultimately, data analysis is a personal choice, and researchers would do well to explore the options that are available. That said, *PyPlr* includes a set of pandas-reliant scripting tools for implementing a standard data processing pipeline that is optimised to study the PLR and to account for some of the idiosyncrasies of Pupil Labs data. These tools are organised into three separate modules: *pyplr.utils* has tools for loading data and extracting trials; *pyplr.preproc* has tools for masking, interpolating and filtering pupil data; and *pyplr.plr* supports pupillometer-style plotting and parametrisation of PLR data. These tools are in continuous development and will evolve over time, hopefully with contributions from other active researchers.

## Examples

We offer two example applications of *PyPlr* and our own custom-built stimulation and measurement system. In the first example, we obtain repeated measurements of the PLR to white light stimuli and compare the results with those from an industry-leading automated pupillometer. In the second, we measure the PIPR to long and short wavelength light.

### Simple PLR

Automated pupillometers are the standard instruments for measuring the PLR. These handheld devices are aimed at the eye to deliver a light stimulus and use infrared video recording and internal algorithms to provide an instant readout of the PLR and its associated parameters. The PLR-3000 (NeurOptics, Laguna Hills, CA, USA) is a leading example with established intraoperator reproducibility and normative benchmarks (Asakawa & Ishikawa, 2017; Winston et al., 2019), access to raw data, and the flexibility to define stimulation protocols by adjusting the pulse intensity, background intensity, measurement duration, pulse duration and pulse onset time. Our system is no competition for the compactness, portability and ease of use of an automated pupillometer like the PLR-3000, but here we demonstrate how it can be made to function in a similar way and to yield comparable results.

### Method

#### Participants

Three non-naive subjects took part in this study, which was approved by The University of Oxford’s central research ethics committee (R54409/RE005). All participants had normal colour vision, as assessed by The New Richmond HRR Pseudoisochromatic Test for Colour Vision (Cole et al., 2006).

#### Stimulation protocols

A PLR-3000 (NeurOptics, Laguna Hills, CA, USA) automated pupillometer was configured to record nine seconds of data and to deliver a one second pulse (180 uW setting) against a dark background after one second of recording. A comparable stimulus for STLAB was generated by obtaining spectral measurements of the PLR-3000 stimulus—produced by four white LEDs—with an OceanOptics STS-VIS (Ocean Insight Inc., Oxford, UK) spectrometer at the usual eye position and then using linear algebra to find the STLAB settings required to produce a spectrum matched for *a*-opic (S-cone-opic, M-cone-opic, L-cone-opic, rhodopic and melanopic) irradiance (CIE, 2018: see Figure 10). The optimal settings were then used to make a one second pulse stimulus for STLAB, which was administered from a Windows laptop running Pupil Capture (v3.2-20) and a custom Python script designed to mimic the functionality of the PLR-3000. Pupil Core Eye camera resolution was kept at (192, 192) with Absolute Exposure Time of 25, and the corresponding settings for the World camera were (640, 480) and 60. Auto Exposure Mode was set to ‘manual mode’ for all cameras, and Auto Exposure Priority was disabled for the World camera.

**Figure 10.**
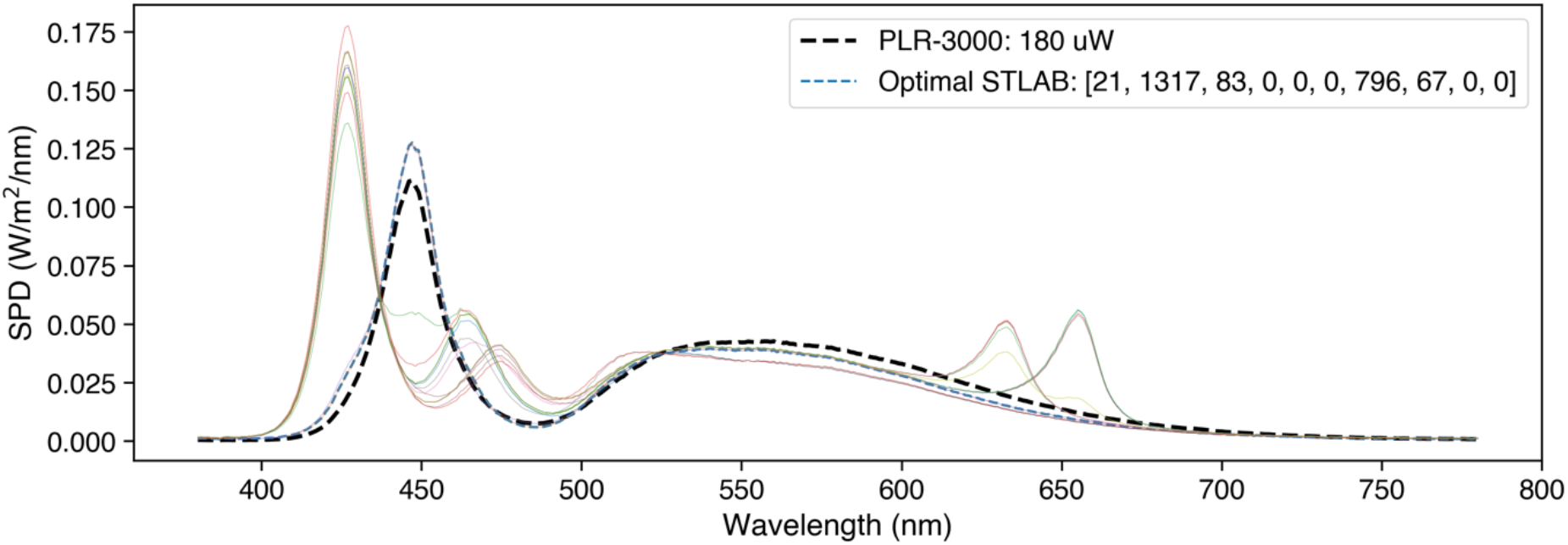
Spectral power distributions of the PLR-3000 white light stimulus and *a*-opic irradiance-matched STLAB-sphere stimulus. We defined the optimal settings as those which produce a spectrum with the least squared error, although in theory it should not matter which is used. Colored lines show alternative solutions to the stimulus matching problem.

To give further insight into the performance of both systems at simple PLR measurement we collected additional data from Subject 1 with different intensity light stimuli. In this comparison, PLR measurements (*n* = 5) were obtained for each of the five stimulus intensity settings on the PLR-3000 (1, 10, 50, 121, 180 uW) and with our own system using theoretically matched stimuli. This time, stimuli were matched using an unconstrained local optimisation procedure (i.e., SciPy’s optimise.minimise function with the ‘SLSQP’ solver: Virtanen et al., 2020) that sought to minimise the difference in *a*-opic irradiance between the measured spectrum for the PLR-3000 180 uW setting and the predicted spectrum for STLAB’s 12-bit LED settings, assuming a linear relationship between input power and radiant flux for both devices. The resulting stimuli were closely matched in terms of their spectral power distributions and *a*-opic irradiances (Figure 12), though they differed slightly with respect to chromaticity due to the mixing of primaries with STLAB.

**Figure 11.**
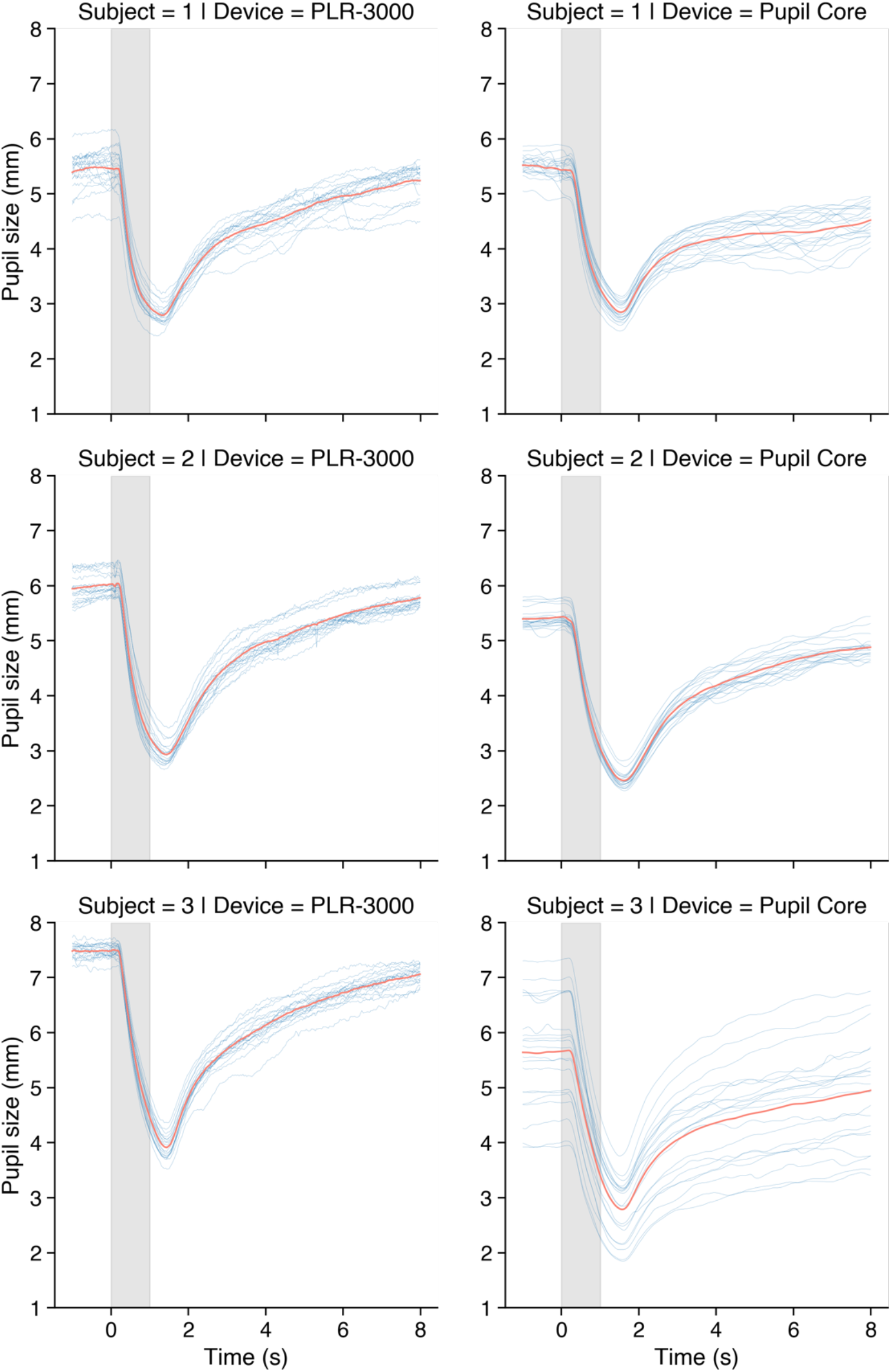
Comparison of PLR measurements (*n* = 20) obtained with a PLR-3000 (180 uW setting) and our own stimulation and measurement system (matched stimulus). The variability in absolute units (i.e., pupil size in millimetres) for Subject 3’s Pupil Core traces was caused by inconsistencies in 3D model fitting and camera repositioning between measurements, which were necessary for optimal pupil tracking. Pupil Core data were filtered with a 3^rd^ order Butterworth filter (4 Hz cut-off).

**Figure 12.**
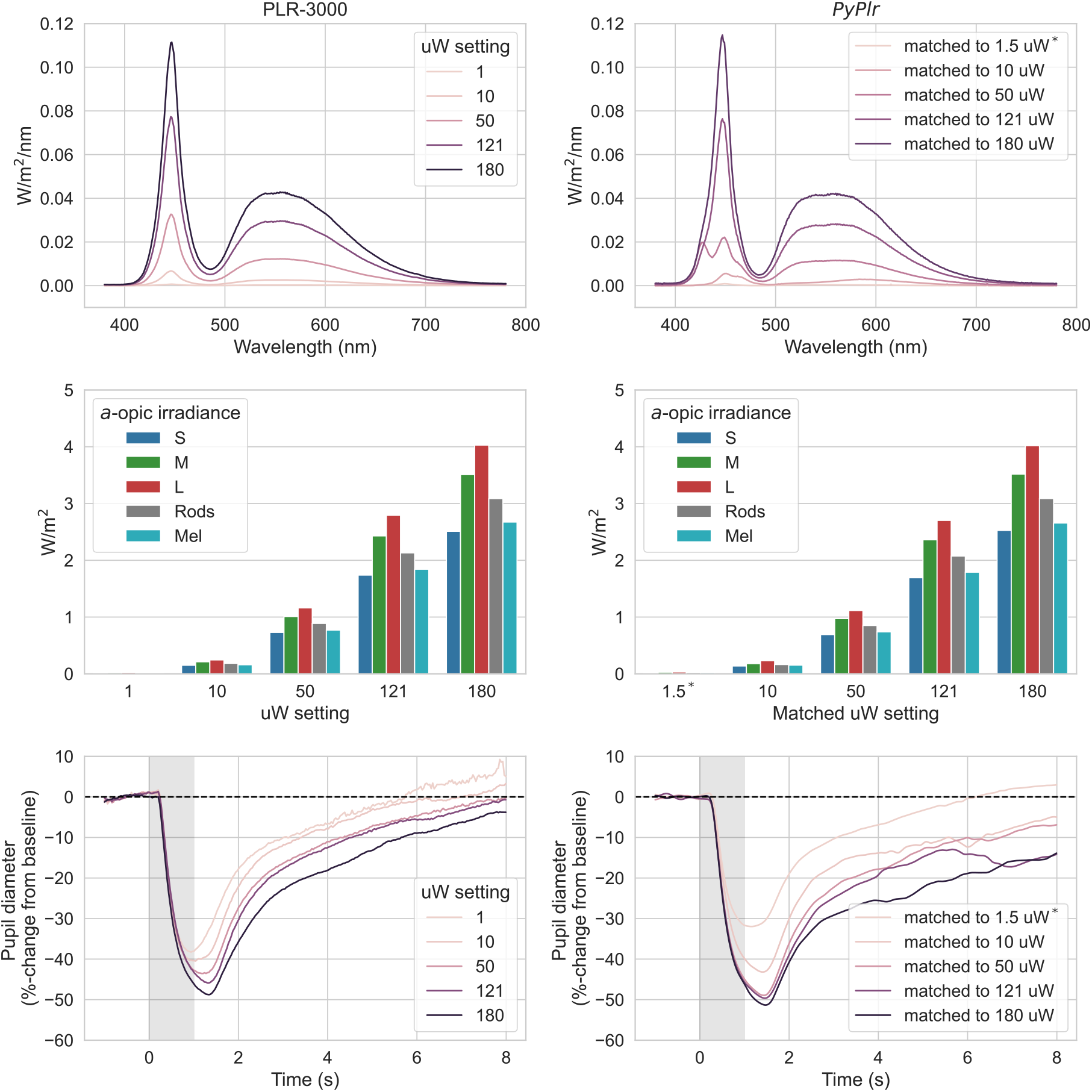
Comparison of PLR measurements to different intensity light stimuli with both systems (left: PLR-3000, right: *PyPlr*). The stimuli were well-matched in terms of spectral power distributions (top row) and *a*-opic irradiances (middle row), though there were slight differences in chromaticity due to the mixing of primaries with STLAB. Average PLR traces (obtained from Subject 1, *n* = 5) for each stimulus intensity (bottom row) followed the expected pattern. Pupil Core data were filtered with a 3^rd^ order Butterworth filter (4 Hz cut-off). ^*^The *.light_stamper(…)* method could not detect the light in the integrating sphere for the 1 uW stimulus match (13.9 lux). We therefore present data for a 1.5× scaled version of the stimulus (20.7 lux), which was detected reliably.

#### Testing procedure

Testing took place in a dark room where the light from the computer monitor was the only source of illumination. PLRs were measured alternately with each system. PLR-3000 measurements were obtained from the right eye and following the manufacturer’s standard guidelines. Pupil Core measurements were obtained from the left eye. For the STLAB-sphere PLRs, eye level was maintained with a chinrest at the vertical centre of the viewing port and an eye patch was worn over the right eye to ensure dose equivalence with the monocular PLR-3000 stimulus. Participants were asked to look straight ahead, to maintain steady fixation, and to refrain from blinking during the recording. If poor results were obtained for any measurement with either system, the measurement was repeated after a short break.

#### Data analysis

PLR-3000 data were obtained from the device via Bluetooth, converted to CSV format and then processed with custom Python software. Invalid samples (marked as 0 in the data file) were masked and reconstructed with linear interpolation. Our custom *PyPlr* application collected data in real-time using the *.light_stamper(…)* and *.pupil_grabber(…)* methods. High frequency noise was removed with a 3^rd^ order Butterworth filter (4 Hz cut-off) before parameters were calculated with *pyplr.plr.PLR*. Raw data and parameters for each measurement were saved in CSV format. Sometimes the pupil failed to reach 75% recovery within the measurement period (see Table 1). In these cases the value of the relevant parameter was treated as ‘not-a-number’ in the averaging procedure.

**Table 1.**
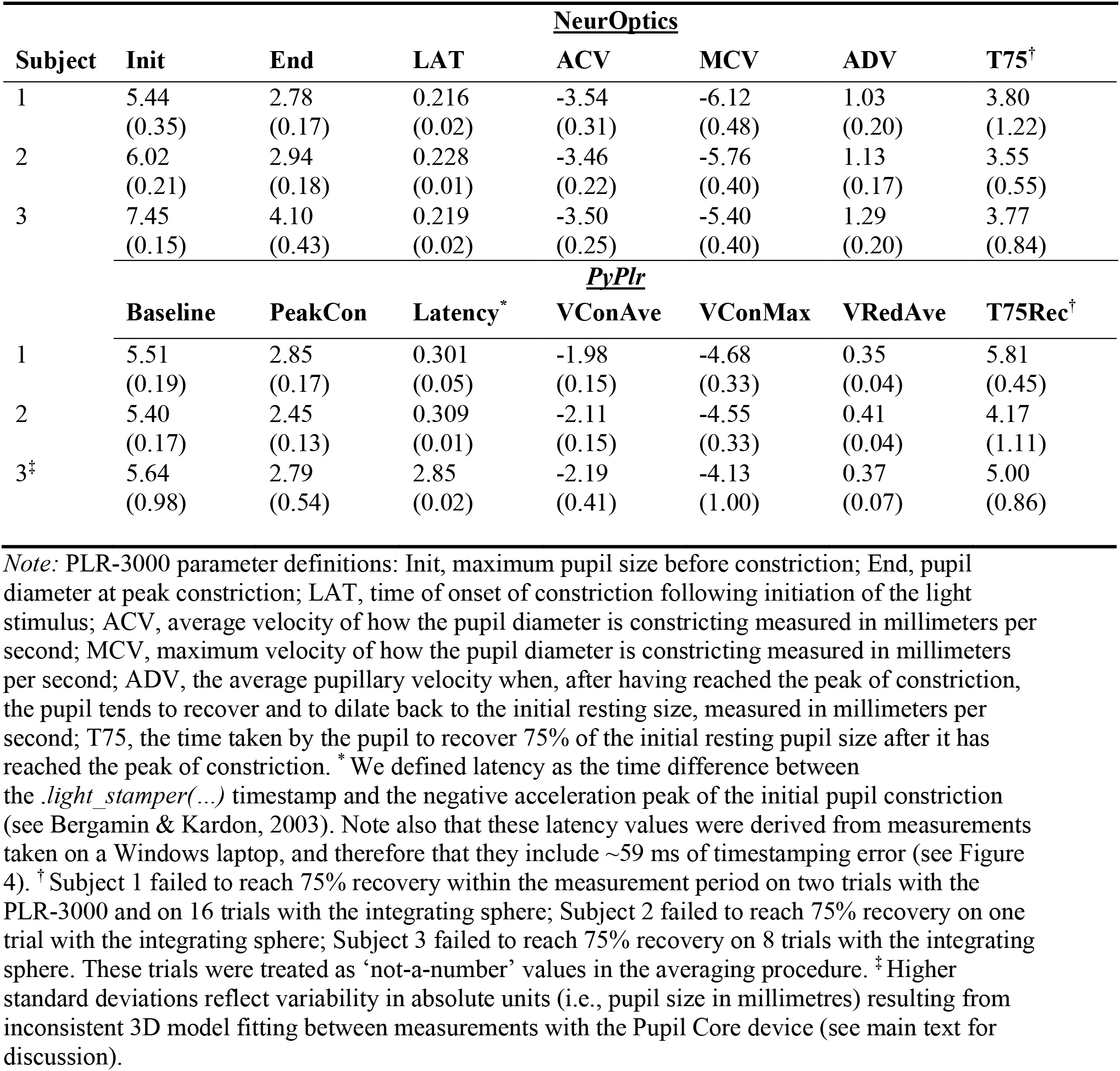
Mean and standard deviation of PLR (*n* = 20) parameters as calculated by a NeurOptics PLR-3000 (NeurOptics, Laguna Hills, CA, USA) and our own system. Different naming conventions emphasize that our parameter calculation principles may differ to those used by the PLR-3000. Note that *pyplr.plr.PLR* calculates other parameters as well.

#### Results

The PLR measurements (*n* = 20) obtained with the PLR-3000 (180 uW setting) and with our own system (theoretically matched stimulus) were comparable in shape and magnitude for all subjects (Figure 11). In terms of absolute units (i.e., pupil size in millimetres), Subjects 1 and 2 showed a close correspondence between devices whereas the data for Subject 3 were more variable due to difficulties in obtaining a consistent 3D model fit between measurements (see general discussion). Discounting the effect of variability in absolute measurement units for Subject 3, the PLR parameters calculated for both systems, shown in Table 1, were also generally comparable.

The additional PLRs collected from Subject 1 using different intensity light stimuli with both systems also followed the expected pattern (Figure 12, bottom row). Differences in the overall shape and magnitude of the PLR traces may reflect stimulus geometry and how the data were processed. Note that we were unable to obtain PLRs with the 1 uW stimulus match as the light was very dim (13.9 lux) in the integrating sphere and could not be detected by the *.light_stamper(…).* As an alternative we present data from a 1.5× scaled version of the stimulus (20.7 lux), which was detected reliably.

#### Discussion

Here we show that our *PyPlr* stimulation and measurement system can function like an industry-leading automated pupillometer. Both systems were configured to record nine seconds of data and to deliver one-second pulses of light stimuli matched for *a*-opic-irradiance (Figure 10). The shape and magnitude of the resulting PLR traces were highly comparable between systems, though there was some variability in terms of absolute units for Subject 3’s PLRs due to difficulties in getting a consistent 3D model fit between measurements (see general discussion).

The PLR-3000 device yields seven parameters for every measured pupil trace, an aspect of functionality that we were able to mimic with *pyplr.plr.PLR* (see Table 1). Despite alternative approaches to calculating the parameters, the averages and standard deviations were generally similar. The most marked discrepancies were between the parameters representing constriction latency and the time taken for the pupil to recover to 75% of the baseline value after reaching peak constriction. Regarding latency, we note that the *PyPlr* data were collected on a Windows laptop and therefore that they include on average ~59 ms timestamping error (corresponding to the average difference between the World and Eye camera timestamps in Figure 4 for OS = Windows | FPS = 120). Subtracting 59 ms from the averages for Subjects 1, 2 and 3 gives values of 242 ms, 250 ms and 226 ms, respectively, which are more plausible with respect to normative values in the literature (e.g., Shah et al., 2020; Straub et al., 1992; Winston et al., 2019). For the 75% metric, the discrepancy may be explained by geometrical differences in retinal stimulation: Although the stimuli were matched for *a*-opic irradiance and delivered monocularly, the PLR-3000 light stimulus comes from 4 small LEDs positioned close to the eye, whereas our integrating sphere system stimulates the entire visual field with reflected light. This may have altered the extent to which the pupil response was driven by melanopsin excitation, which in turn could explain why Subject 1 failed to reach 75% recovery on 16/20 trials with the sphere but only 2/20 trials with the PLR-3000.

Although subtracting 59 ms from our latency measures gives plausible values, we do not advocate for this as a blanket solution. Rather, we point out that the ground truth for constriction latency is difficult to obtain and that measurements are constrained by hardware and calculation principles. For example, with video recording at 30 and 120 frames per second, precision is limited to 33.333 ms and 8.333 ms, respectively, though this could be improved by upsampling the data prior to calculation (e.g., see Bergamin & Kardon, 2003). Similarly, latency measures based on the negative acceleration peak of pupil constriction (e.g., Bergamin & Kardon, 2003) will differ from those based on the time taken to cross a threshold of change from baseline (e.g., Maynard et al., 2015). Repeatability is what ultimately matters in this domain, and our data suggest that both the PLR-3000 (NeurOptics, Laguna Hills, CA, USA) and Pupil Core (Pupil Labs GmbH, Berlin, Germany) systems perform well in this regard.

As expected, PLR measurements showed a graded response to different intensity light stimuli for both devices. Divergences in the shape and magnitude of the pupil traces for each of the stimulus levels may reflect imperfect stimulus matching (e.g., due to the possibly flawed assumption of linearity) and differences in stimulus geometry. It is noteworthy that the *.light_stamper(…)* was unable to detect the light for the 1 uW (13.9 lux) stimulus match in the integrating sphere, even with a detection threshold value of one. This indicates that the *.light_stamper(…)* method may be unsuitable for timestamping very subtle illuminance increments under certain conditions.

### PIPR

Whereas the PLR refers to the general response of the pupil to light, the PIPR describes the sustained constriction of the pupil following exposure to short-wavelength (blue) light and is assumed to be a unique non-invasive signature of melanopsin processing in the human retina. As an optimum protocol for measuring the PIPR, Park et al. (2011) recommend comparing pupil responses to high intensity (2.6 log cd/m^2^) one-second pulses of short and long wavelength light presented in darkness following a period of dark adaptation. Park et al. obtained their PIPR measurements using the industry-leading Espion V5 system with ColorDome LED full-field stimulator (Diagnosys LLC, Lowell, MA). Here we describe a comparable protocol for measuring the PIPR with our own stimulation and measurement system.

### Method

#### Participants

The same participants as previous took part in this study.

#### Stimulation protocols

Stimuli were administered via STLAB’s fourth (blue, λ-max = 470) and tenth (deep red, λ-max = 657) LED channels, which offer maximal and minimal melanopic excitation, respectively. The blue stimulus was set at ~800 lx and the red stimulus was matched for unweighted irradiance. The spectral power distributions of the stimuli are visualised in Figure 13 along with the spectral sensitivity curve for melanopsin. Both were presented for one second using STLAB video files.

**Figure 13.**
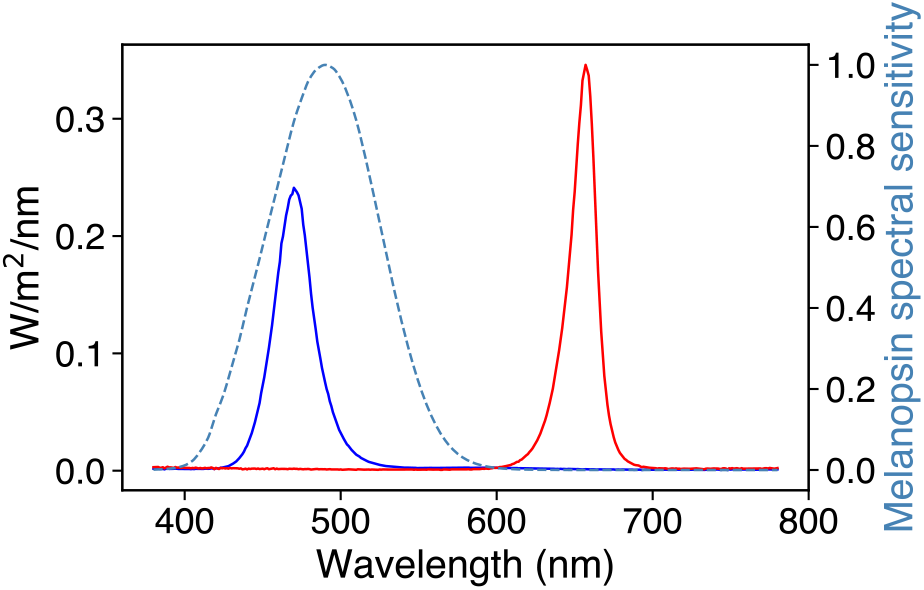
Spectral power distributions of PIPR stimuli shown in relation to the relative energy spectral sensitivity curve for melanopsin.

#### Testing procedure

Participants completed the PIPR protocol in a dark room after 20 minutes dark-adaptation. When ready to begin, they placed their chin on the chinrest and the experimenter ensured that their eyes were level with the vertical centre of the viewing port. Participants were asked to roll their eyes as the experimenter ensured a good fit of the 3D eye models in the Pupil Capture and were then asked to look straight ahead for the duration of the recording. The recording lasted ~12 min, during which time three of each colour light stimulus were administered in a random order with ~2 minutes spacing. A high-pitched beep signalled to the participant that a stimulus would be presented in the next five to ten seconds (in a time-jittered fashion to avoid expectancy effects), and a low-pitched beep indicated that one minute had passed since the stimulus. Recording was binocular at 120 Hz and light stimuli were timestamped using the *.light_stamper(…)* method.

#### Data analysis

Data were exported to CSV format via the Pupil Player software and processed with scripting tools from *pyplr.utils* and *pyplr.preproc*. For each participant, the eye with the highest average confidence was chosen for analysis. To account for blinks, pupil data were masked with ‘not-a-number’ values where the first derivative exceeded ± 3 SD or if the corresponding confidence value was below.95. The missing data were reconstructed with linear interpolation before the time-course was smoothed with a third-order Butterworth filter (4 Hz cut-off). Relative to the *.light_stamper(…)* timestamps, 65 seconds of pupil data were then extracted for each stimulus event and converted to %-change from the average pupil size in a 5 seconds prestimulus baseline.

#### Results

Clear PIPRs were observed for all subjects (Figure 14). Subject 2 tended to blink more often at stimulus onset, which may explain the reduced PIPR to blue stimuli.

**Figure 14.**
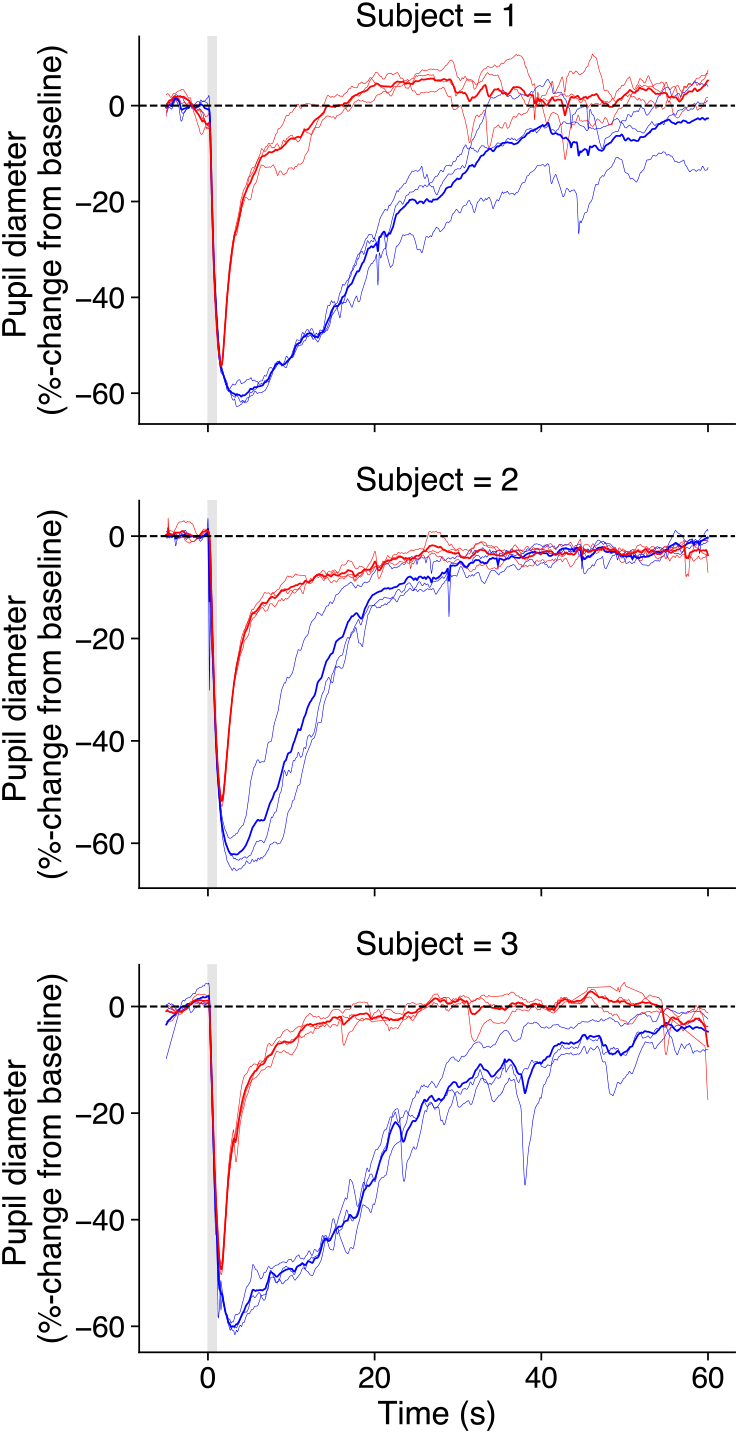
Average PIPRs for Subject 1 (left) and Subject 2 (right). Desaturated lines show individual trials.

#### Discussion

Here we show that our system of hardware and software can be used to measure the PIPR in a way that compares to industry-leading commercial equipment (e.g., Lei et al., 2014; Park et al., 2011; Romagnoli et al., 2020). It is worth noting that many aspects of this protocol are customisable. For example, the duration, intensity, and spectral composition of the stimulus can be specified in accordance with the limits imposed by STLAB. Further, rather than administering simple light pulses, one could generate time-varying stimuli (e.g., sinusoidal flicker). Such stimuli have been used previously to probe the temporal characteristics of melanopsin’s and other photoreceptor’s contributions to pupil control (e.g., Joyce et al., 2015, 2018; Maynard et al., 2015; Rukmini et al., 2019; Spitschan et al., 2014).

## General discussion

In this paper we have described *PyPlr* (Martin & Spitschan, 2021)—a *pip* installable Python software for researching the PLR with the Pupil Core eye-tracking platform. A key feature of *PyPlr* is its feature-rich, object-oriented interface to Pupil Core which includes a *.light_stamper(…)* method for accurate timestamping of any light stimulus (> ~20 lux) given a suitable geometry, and a *.pupil_grabber(…)* method which simplifies real-time access to pupil data. The *.light_stamper(…)* works flawlessly with our own integrated system for a range of practical intensities and we can confirm that it also works with other light sources, such as a computer monitor controlled by *PsychoPy* and a light switch in a dark room (see online documentation for examples). *PyPlr* also has native support for our chosen light stimulation and measurement hardware—STLAB and OceanOptics STS-VIS—as well as tools for streamlining the processing and analysis of pupillometry data. As such, *PyPlr* in combination with Pupil Core is a versatile, extensible and comparatively affordable solution to researching the PLR.

In addition to the software, we have described a low-cost integrating sphere stimulation rig that delivers full field, “Ganzfeld” light stimulation. The integrating sphere provides good control over the geometry of retinal stimulation without the need for a complex Maxwellian view optical setup. The raw materials for our sphere cost us less than £1500, which is a small fraction of the price of an equivalent commercial solution. We use our sphere with an STLAB light engine, giving us a high level of control over the temporal and spectral properties of light stimuli; and we calibrated the system with an OceanOptics STS-VIS spectrometer placed at the normal eye position. Prospective users may wish to develop for alternative stimulation and measurement hardware, in which case their contributions to the software would be greatly appreciated.

We gave two examples showing how our complete integrated setup can rival industry leading commercial equipment for measuring the PLR and PIPR. In the main PLR example, our system was made to function like an automated pupillometer, administering a flash of white light and saving raw data, a plot, and parameters of the PLR. In terms of absolute units and variability, the PLR measurements and parameters were generally comparable to those obtained with an industry-leading automated pupillometer (PLR-3000, NeurOptics, Laguna Hills, CA, USA) under the same stimulus and testing conditions (but see caveat below). Likewise, we were able to obtain measurements of the PIPR which rival those made with industry-leading commercial equipment (e.g., Lei et al., 2014; Park et al., 2011; Romagnoli et al., 2020). Of note, these two examples represent only a snapshot of our system’s capabilities, and the scope for further stimulation and measurement protocols is limited only by the capabilities of Pupil Core and the chosen light source. For example, with STLAB’s 10 LED channels, one could potentially design protocols that use the method of silent substitution to examine the contribution of individual photoreceptor classes to pupil control (e.g., see Spitschan & Woelders, 2018).

The PLRs for Subject 3 (Figure 11, bottom row) and the additional PLRs collected for Subject 1 with different intensity light stimuli (Figure 12, bottom row) highlight important issues that researchers should consider before investing in equipment. First, the issue of absolute units. Pupil Core’s *diameter_3d* data (i.e., pupil size in millimetres) are derived from a pupil detection algorithm that implements a mathematical 3D eye model (Dierkes et al., 2018, 2019; Świrski & Dodgson, 2013). These data have the advantage of being robust to the effects of pupil foreshortening (e.g., see Hayes & Petrov, 2015), but inaccuracies and inconsistencies can still arise from implicit model assumptions, camera positioning and software settings. Such was the case with Subject 3, for whom there were numerous pupil detection issues necessitating camera adjustments and model refits between measurements. This issue may not pose a problem for research applications where the focus is on %-change from baseline, but if researchers are interested in obtaining consistent measurements of absolute pupil size, then an alternative device such as the PLR-3000 may be more suitable. Second, the *.light_stamper(…)* was unable to detect small illuminance increments (< ~20 lux) in our integrating sphere, meaning it may be unsuitable for low light applications under certain stimulus geometries. In such cases, an alternative timestamping protocol may be required.

## Summary

*PyPlr* and Pupil Core offer an affordable, flexible, research-grade solution for studying the PLR. We hope that other researchers will find it useful and contribute to its development.

## Acknowledgements

We thank Grahame Faulkner and David Sliney for guidance on light safety, Chris Gibbs, Chris Hatcher, David Ilsley and Duncan Constable for building the stimulation rig, Thibault Lestang and the Oxford Research Software Engineering (OxRSE) Group for helpful advice at various stages of software development, and Pablo Prietz and the Pupil Labs community for support on Discord.

## Open Practices Statement

The data and materials for all experiments are available at https://zenodo.org/record/4785288. None of the experiments were preregistered.

